# Coleoborers (Curculionidae: Scolytinae) in native and homogeneous systems of Brazil nut (*Bertholletia excelsa* bonpl.) in the Southern Amazon, Brazil

**DOI:** 10.1101/2020.05.26.116319

**Authors:** Marcus Henrique Martins e Silva, Juliana Garlet, Fernando Luis Silva, Carla da Silva Paula

## Abstract

Brazil nut is one of the most important species of the Amazon due to its socioeconomic importance. Especially in homogeneous production systems, it may be susceptible to damage by wood-boring insects, as by the subfamily Scolytinae (Coleoptera: Curculionidae); thus, inadequate management conditions can cause economic damage. Therefore, the objective of the present work is to evaluate the occurrence of wood-boring insects (Curculionidae: Scolytinae) in native and homogeneous systems of Brazil nut in the Meridional Amazonian, Brazil. The study was conducted in three environments: Conserved Native Planting nut, Anthropized Native Planting nut and Homogeneous Planting nut. Twelve ethanol traps were installed in each environment during four sampling periods. The data were submitted to entomofaunistic analysis, Pearson’s correlation analysis and cluster analysis. A total of 2,243 individuals from 31 species were sampled, of which 23 were from the Anthropized Native Planting nut, 24 from the Homogeneous Planting nut and 26 from the Conserved Native Planting nut. In the faunistic analysis, we highlight the species *Xyleborus affinis* (Eichhoff, 1868), which was the most representative one in the three environments and a super-dominant species in all four sampling periods. There was a greater similarity between the Anthropized Native Planting nut and the Conserved Native Planting nut; these two environments showed dissimilarity with the Homogeneous Planting nut. Monitoring coleoborers in Brazil nut agroecosystems is fundamental for the establishment of integrated pest management strategies.

## Introduction

Considered one of the most important extractive species in the Amazon and Brazil, Brazil nut (*Bertholletia excelsa* Bonpl.: Lecythidaceae) is part of the socioeconomic support base of many traditional communities and has become a crop of interest for commercial exploitation in homogeneous forest stands [1–3].

Mainly because of deforestation, the decrease in Brazil nut trees in natural areas compromises the sustainability of the extractive production chain. It is possible to point to a tendency for declining extractive activity and, at the same time, the potential for the rise of Brazil nut plantations, with greater technification, enhanced management strategies and the use of sustainable agricultural practices [4,5]. In this way, the development of silvicultural studies of Brazil nut as an alternative in the exploration of forest and non-forest products is of paramount importance, aiming at the development of effective management strategies [6].

In forest systems, insects perform fundamental functions and interactions in maintaining and regulating the conditions and resources of ecosystems. However, insect-plant interactions in certain circumstances can compromise production in agroecosystems, whether in direct or indirect damage, due to log boring, defoliation, seed drilling in the field or in storage; they may also be vectors of several plant diseases. Attacks of *Hypothenemus hampei* (Ferrari, 1867) have been verified in Brazil nut stands in southeastern Pará, making this coleoborer a potential cause of damage in these production systems [7]. In addition, *Tribolium castaneum* (Herbst, 1797), *Rhyzopertha dominica* (Fabricius, 1792), *Ephestia kuehniella* (Zeller) and *Plodia interpunctella* (Hübner) have great potential to cause damage to stored almonds [8–12].

Forest massifs, characterised by homogeneous planting systems, become susceptible to the development of insect-plant interactions harmful to crops, especially when considering the Amazon region, which has a rich biodiversity and potential for the emergence of new interactions of this type [13]. The implementation of homogeneous plantations can cause an increase in insects harmful to forest species, making it necessary to conduct population studies of possible pests to outline strategies to reduce negative impacts, especially in the Southern Amazon, where studies of wood-boring insects are still scarce [14].

Wood-boring insects, in particular those belonging to the subfamily Scolytinae (Coleoptera: Curculionidae), form one of the most important groups of forest pests [15], with more than 6,000 described species [16,17]. This group of insects has great potential for damage by promoting the opening of galleries in the tree trunks making them weak and stressed, in addition to allowing the infection of plant tissues by fungi [18].

To understand the events involving the occurrence of wood-boring insects, one must primarily consider the proper identification of the insect [16]; thus, a monitoring program becomes essential to assess population levels. In this context, the objective of the present work is to evaluate the occurrence of wood-boring insects (Scolytinae) in three different production systems of Brazil nut (*Bertholletia excelsa* Bonpl.) in the Southern Amazon.

## Materials and Methods

### Study area

The study was developed in three Brazil nut areas in the Southern Amazon, characterised by different compositions, biophysical and ecological gradients, management levels and anthropogenic factors: Brazil Nut Native Anthropized (Environment 1), inserted in a fragment of native forest and pasture of *Panicum maximum* Jacq. (Poaceae), located in Alta Floresta, Mato Grosso, with a total area of 22 ha. Homogeneous Brazil Nut (Environment 2), located in Paranaíta, Mato Grosso, has a total area of 28 ha, spaced 6 x 6 m, with an age of approximately 17 years. This system presents a high level of densification, since there has been no thinning since its implantation, which has promoted a greater overlap of crowns and increased shading. Brazil Nut Native Preserved (Environment 3) is inserted in an area of dense forest and covers 250 ha, with a significant level of conservation, located at the Experimental Station of the Executive Committee of the Plan of the Cacao Plantation in Alta Floresta.

The region’s climate is classified as Aw, with bimodal precipitation and a clear distinction between two seasons, a dry winter and a rainy season [19]. The annual average temperature is 26°C, with an annual precipitation between 2,800 and 3,100 mm, concentrated between November and May.

### Sampling and identification of species

To monitor the occurrence of insects in the study areas, 36 ethanolic traps (attractive alcohol 96° GL) were used, adapted from the Pet-Santa Maria model [20] and installed at a height of 1.5 m. In each study area, 12 traps were established at three sampling points, which were composed of a set of four traps arranged in the shape of a cross and 40 m apart.

Based on the characteristics of rainfall distribution in the Alta Floresta region [21], four sampling periods were established between the years 2018 and 2019: 1st period - October/November (beginning of the rainy season), 2nd period - January/February (full rainy season), 3rd period - May (beginning of the dry season), 4th period - August (full dry season). The traps remained operative in the field for 22 days in each of the four periods, with collections performed at intervals of seven days.

After collection in the field, the insects were screened according to their morphological characteristics and quantified considering the study environments and sampling period. Subsequently, the individuals were dried in an oven at 38° for 24 h and sent for identification at the species level to Dr. Eli Nunes Marques of the Federal University of Paraná, Brazil.

Climatological data of relative humidity, temperature and precipitation were analysed with the records of the Surface Meteorological Station of the National Institute of Meteorology in Alta Floresta, considering the 90 days prior to the date of the last collection of each sampling period.

### Data analysis

The studied environments were compared based on the diversity indices, entomofaunistic variables and similarities. Entomofaunistic analysis was performed with the ANAFAU [22] software, which classifies the species according to their dominance, abundance, frequency and constancy. To assess the similarity between environments, the Bray-Curtis grouping and the UPGMA distance algorithm were adopted. The diversity of the environments was calculated using the Shannon-Wiener (H’) and Simpson (Ds) indices and Equitability through the Pielou index (J). The climatic data for each sampling period were subjected to Pearson’s correlation analysis (r) (p ≤ 0.05), with the abundance values of the species considered ecological indicators (species that reached the maximum category in each analysed fauna variable) in the entomofaunistic analysis to understand the seasonality of each species as a function of variations over the four sampling times. For elaboration of the statistical and diversity analyses, the PAST statistical software was used [23].

## Results

Over the four sampling periods, 2,243 individuals were collected, of which 517 (23.05%) occurred in Environment 1, 1,233 (54.97%) in Environment 2 and 493 (21.98%) in Environment 3. In relation to species richness, among the 31 species identified, 23 were found in Environment 1, 24 in Environment 2 and 26 in Environment 3. Table 1 shows the entomofaunistic analysis of the species of Scolytinae in the three different environments.

In Environment 1, 10 species were considered dominant or super-dominant, accounting for 43% of the species. Environment 2 was characterised by 11 dominant or super-dominant species, equivalent to 45% of the total species. In Environment 3, there were 10 dominant or super-dominant species, representing 38% of the total species.

**Table 1.** Entomofaunistic analysis of Scolytinae in the different environments analysed in the Southern Amazon.

| ESPÉCIE | AMBIENTE 1 |  |  |  |  |  | AMBIENTE 2 |  |  |  |  |  | AMBIENTE 3 |  |  |  |  |  |
| --- | --- | --- | --- | --- | --- | --- | --- | --- | --- | --- | --- | --- | --- | --- | --- | --- | --- | --- |
|  | NI | % | D | A | F | C | NI | % | D | A | F | C | NI | % | D | A | F | C |
| *Cnesinus dryografus* Schedl, 1951 | 1 | 0,19 | nd | r | pf | y | 1 | 0,08 | nd | r | pf | y | - | - | - | - | - | - |
| *Coccotrypes palmarum* Eggers, 1933 | - | - | - | - | - | - | 1 | 0,08 | nd | r | pf | y | 7 | 1,42 | d | c | f | w |
| *Corthylus antennarius* Schedl, 1966 | - | - | - | - | - | - | - | - | - | - | - | - | 3 | 0,61 | nd | d | pf | y |
| *Corthylus nudipenis* Schedl, 1950* | 5 | 0,97 | nd | c | f | w | 31 | 2,51 | d | a | mf | w | 4 | 0,81 | nd | c | pf | w |
| *Corthylus pharax* Schedl, 1976 | 7 | 1,35 | d | c | f | y | - | - | - | - | - | - | 12 | 2,43 | d | c | mf | y |
| *Corthylus populans* Eichhoff, 1868 | 2 | 0,39 | nd | d | pf | y | 2 | 0,16 | nd | r | pf | y | 1 | 0,2 | nd | r | pf | y |
| *Corthylocurus vernaculum* Schedl | 1 | 0,19 | nd | r | pf | y | - | - | - | - | - | - | - | - | - | - | - | - |
| *Cryptocarenum diadematus* Eggers, 1937* | 20 | 3,87 | d | ma | mf | w | 62 | 5,03 | d | ma | mf | w | 36 | 7,3 | d | ma | mf | w |
| *Cryptocarenum hevea* Hagedorni, 1912* | 6 | 1,16 | d | c | f | w | 39 | 3,16 | d | ma | mf | w | 2 | 0,41 | nd | r | pf | y |
| *Cryptocarenum seriatus* Eggers, 1933* | 55 | 10,6 | d | ma | mf | w | 113 | 9,16 | d | ma | mf | w | 21 | 4,26 | d | ma | mf | w |
| *Hypothenemus bolivianus* Eggers, 1931 | - | - | - | - | - | - | - | - | - | - | - | - | 3 | 0,61 | nd | d | pf | w |
| *Hypothenemus eruditus* Westwood, 1836 | 8 | 1,55 | d | c | f | w | 9 | 0,73 | d | c | f | w | 6 | 1,22 | d | c | f | y |
| *Hypothenemus seriatus* Eichhoff, 1972 | 1 | 0,19 | nd | r | pf | y | 2 | 0,16 | nd | r | pf | w | 5 | 1,01 | nd | c | f | w |
| *Premnobiatus cavipennis* Eichhoff, 1878* | 36 | 6,96 | d | ma | mf | w | 41 | 3,33 | d | ma | mf | w | 31 | 6,29 | d | ma | mf | w |
| *Sampsonius dampfi* Schedl, 1940* | 6 | 1,16 | d | c | f | w | 19 | 1,54 | d | c | f | w | 28 | 5,68 | d | ma | mf | w |
| *Sampsonius pedrosae* Schenherr, 1994* | 1 | 0,19 | nd | r | pf | y | 1 | 0,08 | nd | r | pf | y | 18 | 3,65 | d | ma | mf | w |
| *Xyleborinus reconditus* Schedl, 1963 | - | - | - | - | - | - | 3 | 0,24 | nd | r | pf | y | 2 | 0,41 | nd | r | pf | y |
| *Xyleborus adelographus* Eichhof, 1867 | 6 | 1,16 | d | c | f | y | - | - | - | - | - | - | 1 | 0,2 | nd | r | pf | y |
| *Xyleborus affinis* Eichhoff, 1868* | 324 | 62,7 | sd | sa | sf | w | 553 | 44,8 | sd | sa | sf | w | 294 | 59,6 | sd | sa | sf | w |
| *Xyleborus biconicus* Eggers, 1928 | 1 | 0,19 | nd | r | pf | y | - | - | - | - | - | - | - | - | - | - | - | - |
| *Xyleborus biseriatus* Schedl, 1963 | 4 | 0,77 | nd | c | f | w | 2 | 0,16 | nd | r | pf | y | 1 | 0,2 | nd | r | pf | y |
| *Xyleborus brasiliensis* Eggers, 1928 | 3 | 0,58 | nd | d | pf | y | 4 | 0,32 | nd | d | pf | w | 1 | 0,2 | nd | r | pf | y |
| *Xyleborus corniculatus* Schedl, 1948 | 2 | 0,39 | nd | d | pf | y | 3 | 0,24 | nd | d | pf | y | 2 | 0,41 | nd | r | pf | w |
| *Xyleborus ferrugineus* Fabricius, 1801* | 23 | 4,45 | d | ma | mf | w | 20 | 1,62 | d | c | f | w | 7 | 1,42 | d | c | f | w |
| *Xyleborus granulicauda* Eggers, 1931 | 2 | 0,39 | nd | d | pf | y | - | - | - | - | - | - | 2 | 0,41 | nd | r | pf | y |
| *Xyleborus spinulosus* Blandford, 1898* | 2 | 0,39 | nd | d | pf | y | 301 | 24,4 | sd | sa | sf | w | 1 | 0,2 | nd | r | pf | y |
| *Xyleborus squamulatus* Eichhoff, 1868 | - | - | - | - | - | - | 18 | 1,46 | d | c | f | y | 1 | 0,2 | nd | r | pf | y |
| *Xyleborus truncatellus* Schedl, 1951 | - | - | - | - | - | - | 3 | 0,24 | nd | d | pf | y | 2 | 0,41 | nd | r | pf | y |
| *Xylosandrus germanus* Blandford, 1894 | - | - | - | - | - | - | 1 | 0,08 | nd | r | pf | y | 2 | 0,41 | nd | r | pf | y |
| *Xyleborus tolimanus* Eggers, 1928 | - | - | - | - | - | - | 3 | 0,24 | nd | d | pf | y | - | - | - | - | - | - |
| *Xylosandrus reconditus* Schedl, 1963 | 1 | 0,19 | nd | r | pf | y | 1 | 0,08 | nd | r | pf | y | - | - | - | - | - | - |
| TOTAL | 517 | 100 |  |  |  |  | 1233 | 100 |  |  |  |  | 493 | 100 |  |  |  |  |
| Índices de Diversidade e Equitabilidade |  | J | Ds | H' |  |  | J | Ds | H' |  |  |  | J | Ds | H' |  |  |  |
|  |  | 0,48 | 0,58 | 1,53 |  |  | 0,55 | 0,72 | 1,75 |  |  |  | 0,52 | 0,62 | 1,71 |  |  |  |

NI = number of individuals. D = Dominance - (d): dominant; (nd) non-dominant. A = Abundance - (ma) very abundant; (a) abundant; (c) common; (d) dispersed; (r) rare. F = Frequency - (mf) very frequent; (f) frequent; (mp) infrequent. C = Constancy - (w) constant; (y) accessory; (z) accidental. J = Pielou's evenness. Ds = Simpson's index. H' = Shannon-Wiener index. \*= species classified as an ecological indicator.

The Venn diagram (Fig. 1) shows that 16 species were common among the three environments.

**Fig 1.**
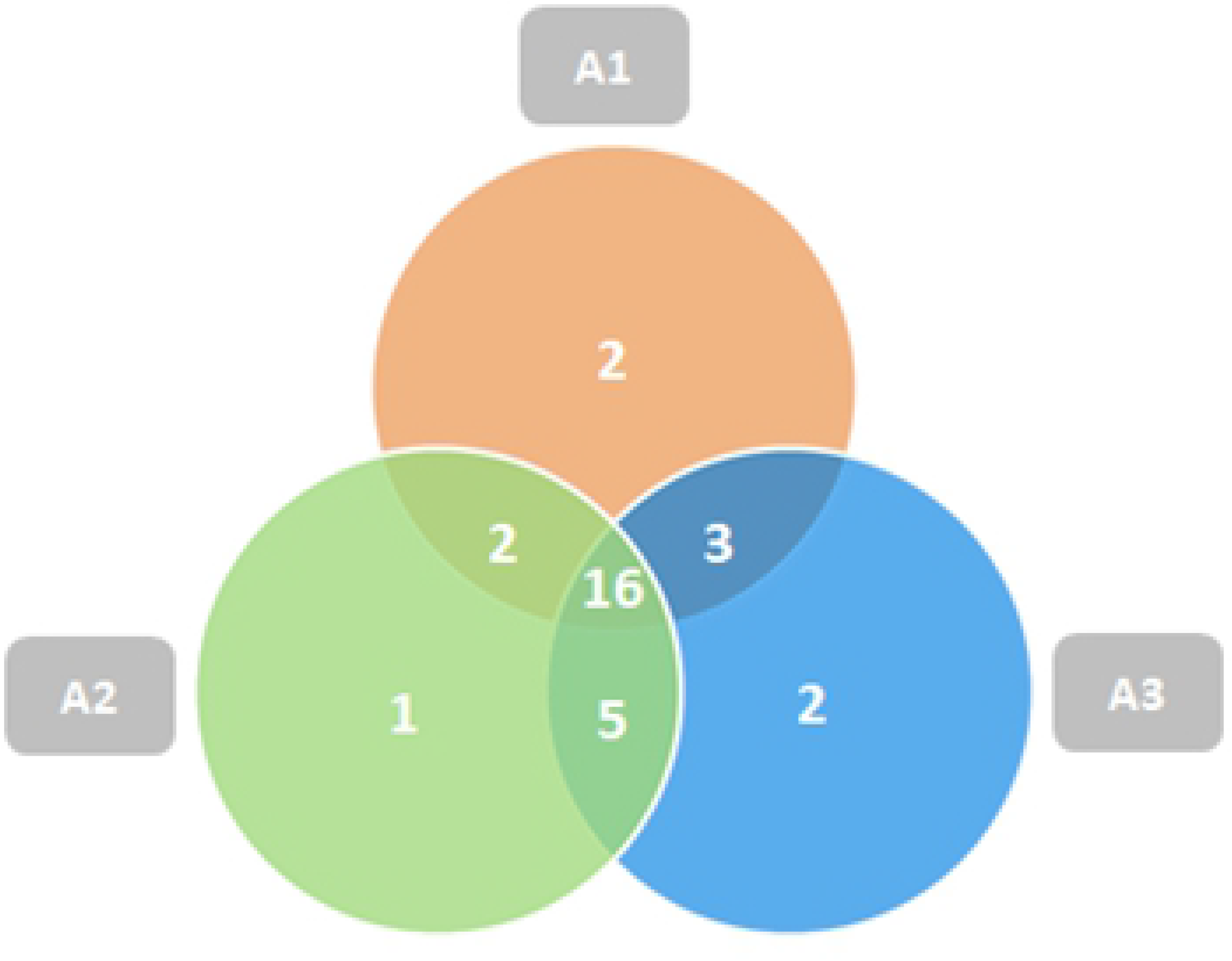
Venn diagram of the distribution of Scolytinae species in different environments for brazil nut cultivation in the southern Amazon.

Some species were restricted by some environments. *Corthylocurus vernaculum* and *Xyleborus biconicus* were verified only in Environment 1, while the species *Xyleborus tolimanus* occurred only in Environment 2. *Corthylus antennarius* and *Hypothenemus bolivianus* were observed only in Environment 3. In general, these species presented frequencies below 1% in their respective environments.

Of the 31 species identified in this study, 9 were classified as ecological indicators, as they reached the maximum categories in all analysed indices, according to the entomofaunistic criteria [22]. Entomofaunistic analyses of these species in the three environments and in each sampling period allow inferring about the effects of environmental conditions on the population density of the Scolytinae and their categories of dominance, abundance, frequency and constancy, as shown in Tables 2–4.

**Table 2.** Entomofaunistic analysis of the Scolytinae indicator species occurring in Environment 1 in four sampling periods in the Southern Amazon.

| ESPÉCIE | PERÍODO 1 |  |  |  | PERÍODO 2 |  |  |  | PERÍODO 3 |  |  |  | PERÍODO 4 |  |  |  |
| --- | --- | --- | --- | --- | --- | --- | --- | --- | --- | --- | --- | --- | --- | --- | --- | --- |
|  | D | A | F | C | D | A | F | C | D | A | F | C | D | A | F | C |
| *C. diadematus* | d | c | f | w | nd | c | f | w | nd | d | pf | w | nd | c | f | w |
| *C. seriatus* | d | ma | mf | w | d | ma | mf | w | nd | c | f | w | nd | c | f | w |
| *P. cavipenis* | d | ma | mf | w | d | c | f | w | nd | d | pf | w | d | ma | mf | w |
| *X. affinis* | sd | sa | sf | w | sd | sa | sf | w | sd | sa | sf | w | sd | sa | sf | w |
| *X. ferrugineus* | d | ma | mf | w | nd | c | f | w | nd | a | mf | w | nd | c | f | w |

NI = number of individuals. D = Dominance - (d): dominant; (nd) non-dominant. A = Abundance - (ma) very abundant; (a) abundant; (c) common; (d) dispersed; (r) rare. F = Frequency - (mf) very frequent; (f) frequent; (mp) infrequent. C = Constancy - (w) constant; (y) accessory; (z) accidental. J = Pielou's evenness. Ds = Simpson's index. H' = Shannon-Wiener index. \* = species classified as an ecological indicator.

In Environment 1, there was variation in the behaviour of the indicator species over the four sampling periods (Table 2). Only the species *Xyleborus affinis* maintained the status of a super-dominant species in the four periods analysed. The species *Cryptocarenus seriatus* behaved as a dominant species in the initial rainy season and at the height of the rainy season. *Premnobius cavipennis* did not present a dominant pattern, except at the end of the rainy season/beginning of the dry season. *Cryptocarenus diadematus* and *Xyleborus ferrugineus* reached the category of dominant species in the initial rainy season.

Analysis of Environment 2 for indicator species (Table 3) showed that the species *Cryptocarenus seriatus*, *Cryptocarenus hevea* and *Xyleborus affinis* maintained their status as dominant or super-dominant species in all sampling periods. *Cryptocarenus diadematus* did not appear to be dominant only in the dry season (period 4). *Premnobius cavipennis* did not present a dominant pattern only in the second period. The species *Xyleborus spinulosus* was classified as dominant and super-dominant in the third and fourth sampling periods, respectively.

**Table 3.**
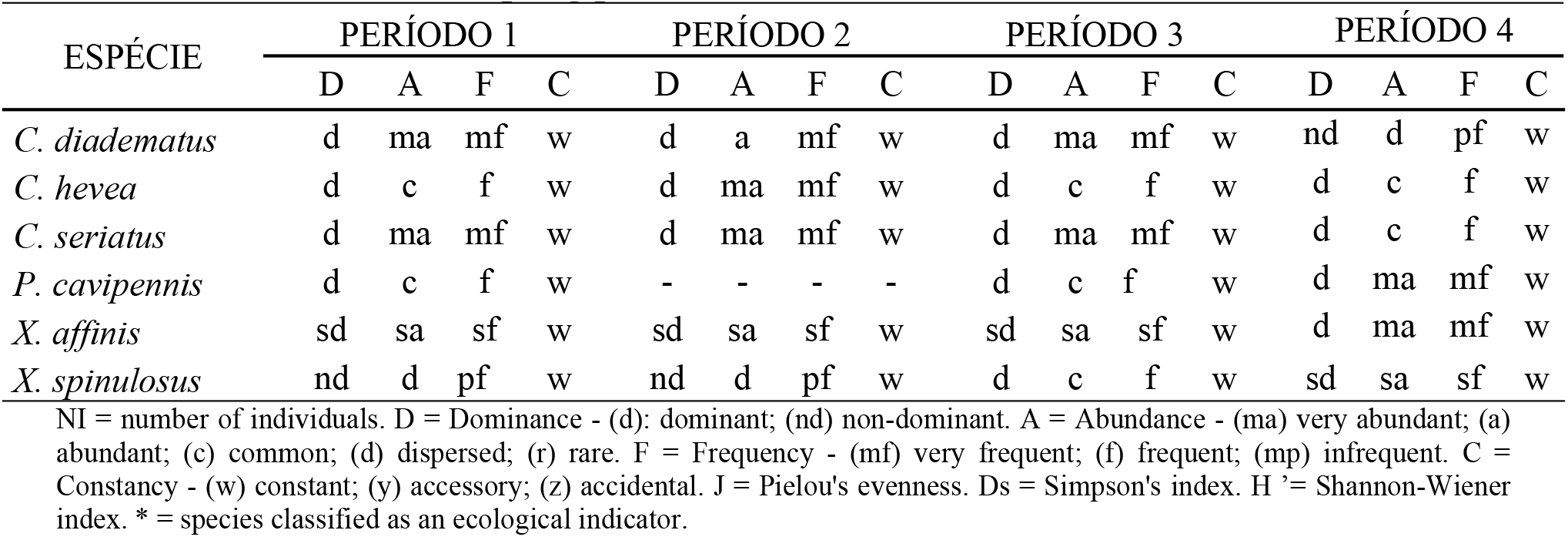
Entomofaunistic analysis of the Scolytinae indicator species occurring in Environment 2 in four sampling periods in the Southern Amazon.

| ESPÉCIE | PERÍODO 1 |  |  |  | PERÍODO 2 |  |  |  | PERÍODO 3 |  |  |  | PERÍODO 4 |  |  |  |
| --- | --- | --- | --- | --- | --- | --- | --- | --- | --- | --- | --- | --- | --- | --- | --- | --- |
|  | D | A | F | C | D | A | F | C | D | A | F | C | D | A | F | C |
| *C. diadematus* | d | ma | mf | w | d | a | mf | w | d | ma | mf | w | nd | d | pf | w |
| *C. hevea* | d | c | f | w | d | ma | mf | w | d | c | f | w | d | c | f | w |
| *C. seriatus* | d | ma | mf | w | d | ma | mf | w | d | ma | mf | w | d | c | f | w |
| *P. cavipennis* | d | c | f | w | - | - | - | - | d | c | f | w | d | ma | mf | w |
| *X. affinis* | sd | sa | sf | w | sd | sa | sf | w | sd | sa | sf | w | d | ma | mf | w |
| *X. spinulosus* | nd | d | pf | w | nd | d | pf | w | d | c | f | w | sd | sa | sf | w |

NI = number of individuals. D = Dominance - (d): dominant; (nd) non-dominant. A = Abundance - (ma) very abundant; (a) abundant; (c) common; (d) dispersed; (r) rare. F = Frequency - (mf) very frequent; (f) frequent; (mp) infrequent. C = Constancy - (w) constant; (y) accessory; (z) accidental. J = Pielou's evenness. Ds = Simpson's index. H' = Shannon-Wiener index. \* = species classified as an ecological indicator.

Table 4 shows that *Xyleborus affinis* and *Cryptocarenus diadematus* maintained their status as dominant species for Environment 3, irrespective of the sampling period. *Sampsonius dampfi* was not classified as dominant only in the second period, while *Sampsonius pedrosae* was classified as dominant in the third period. *Cryptocarenus seriatus* was observed as dominant in the second and fourth periods, and *Premnobius cavipennis* was dominant only in the fourth sampling period.

**Table 4.** Entomofaunistic analysis of the Scolytinae indicator species occurring in Environment 3 in four sampling periods in the Southern Amazon.

| ESPÉCIE | PERÍODO 1 |  |  |  | PERÍODO 2 |  |  |  | PERÍODO 3 |  |  |  | PERÍODO 4 |  |  |  |
| --- | --- | --- | --- | --- | --- | --- | --- | --- | --- | --- | --- | --- | --- | --- | --- | --- |
|  | D | A | F | C | D | A | F | C | D | A | F | C | D | A | F | C |
| *C. diadematus* | d | ma | mf | w | d | ma | mf | w | d | c | f | w | d | c | f | w |
| *C. seriatus* | nd | c | f | w | d | ma | mf | w | - | - | - | - | d | c | f | w |
| *P. cavipennis* | nd | c | f | w | nd | c | f | w | nd | d | pf | w | d | ma | mf | w |
| *S. dampfi* | d | ma | mf | w | nd | c | f | w | d | c | f | w | d | a | mf | w |
| *S. pedrosae* | nd | c | f | w | - | - | - | - | d | ma | mf | w | nd | d | pf | w |
| *X. affinis* | sd | sa | sf | w | sd | sa | sf | w | sd | sa | sf | w | d | ma | mf | w |

NI = number of individuals. D = Dominance - (d): dominant; (nd) non-dominant. A = Abundance - (ma) very abundant; (a) abundant; (c) common; (d) dispersed; (r) rare. F = Frequency - (mf) very frequent; (f) frequent; (mp) infrequent. C = Constancy - (w) constant; (y) accessory; (z) accidental. J = Pielou's evenness. Ds = Simpson's index. H' = Shannon-Wiener index. \* = species classified as an ecological indicator.

In general, the correlation analysis between population fluctuation of species and climatic variables was not statistically significant, as shown in Table 5.

**Table 5.** Analysis of correlations between the population fluctuation of the Scolytinae indicator species and the climatic variables of the different environments in the Southern Amazon.

| Espécies | Tmax | Tmin | Tmed | Umax | Umin | Umed | P(mm) |
| --- | --- | --- | --- | --- | --- | --- | --- |
| Ambiente 1 |  |  |  |  |  |  |  |
| *C. diadematus* | 0,43<sup>NS</sup> | -0,18<sup>NS</sup> | 0,42<sup>NS</sup> | -0,87<sup>NS</sup> | -0,41<sup>NS</sup> | -0,46<sup>NS</sup> | -0,47<sup>NS</sup> |
| *C. seriatus* | -0,11<sup>NS</sup> | 0,26<sup>NS</sup> | 0,58<sup>NS</sup> | -0,77<sup>NS</sup> | 0,08<sup>NS</sup> | 0,01<sup>NS</sup> | 0,04<sup>NS</sup> |
| *P. cavipennis* | 0,76<sup>NS</sup> | -0,76<sup>NS</sup> | -0,28<sup>NS</sup> | -0,89<sup>NS</sup> | -0,84<sup>NS</sup> | -0,88<sup>NS</sup> | -0,85<sup>NS</sup> |
| *X. affinis* | -0,61<sup>NS</sup> | 0,53<sup>NS</sup> | 0,41<sup>NS</sup> | -0,44<sup>NS</sup> | 0,51<sup>NS</sup> | 0,43<sup>NS</sup> | 0,50<sup>NS</sup> |
| *X. ferrugineus* | 0,14<sup>NS</sup> | 0,23<sup>NS</sup> | 0,80<sup>NS</sup> | -0,54<sup>NS</sup> | -0,04<sup>NS</sup> | -0,07<sup>NS</sup> | -0,12<sup>NS</sup> |
| Ambiente 2 |  |  |  |  |  |  |  |
| *C. diadematus* | -0,08<sup>NS</sup> | 0,52<sup>NS</sup> | 0,87<sup>NS</sup> | 0,20<sup>NS</sup> | 0,29<sup>NS</sup> | 0,32<sup>NS</sup> | 0,21<sup>NS</sup> |
| *C. hevea* | 0,07<sup>NS</sup> | 0,36<sup>NS</sup> | 0,89<sup>NS</sup> | -0,32<sup>NS</sup> | 0,08<sup>NS</sup> | 0,06<sup>NS</sup> | -0,01<sup>NS</sup> |
| *C. seriatus* | 0,28<sup>NS</sup> | 0,18<sup>NS</sup> | 0,78<sup>NS</sup> | -0,26<sup>NS</sup> | -0,10<sup>NS</sup> | -0,10<sup>NS</sup> | -0,19<sup>NS</sup> |
| *P. cavipennis* | 0,89<sup>NS</sup> | -0,96* | -0,73<sup>NS</sup> | -0,15<sup>NS</sup> | -0,91<sup>NS</sup> | -0,88<sup>NS</sup> | -0,88<sup>NS</sup> |
| *X. affinis* | -0,47<sup>NS</sup> | 0,64<sup>NS</sup> | 0,81<sup>NS</sup> | -0,43<sup>NS</sup> | 0,48<sup>NS</sup> | 0,42<sup>NS</sup> | 0,43<sup>NS</sup> |
| *X. spinulosus* | 0,71<sup>NS</sup> | -0,94<sup>NS</sup> | -0,92<sup>NS</sup> | -0,13<sup>NS</sup> | -0,80<sup>NS</sup> | -0,79<sup>NS</sup> | -0,75<sup>NS</sup> |
| Ambiente 3 |  |  |  |  |  |  |  |
| *C. diadematus* | -0,16<sup>NS</sup> | 0,32<sup>NS</sup> | 0,64<sup>NS</sup> | -0,73<sup>NS</sup> | 0,13<sup>NS</sup> | 0,07<sup>NS</sup> | 0,09<sup>NS</sup> |
| *C. seriatus* | 0,05<sup>NS</sup> | -0,41<sup>NS</sup> | -0,57<sup>NS</sup> | -0,60<sup>NS</sup> | -0,27<sup>NS</sup> | -0,33<sup>NS</sup> | -0,21<sup>NS</sup> |
| *P. cavipennis* | 0,72<sup>NS</sup> | -0,96* | -0,91<sup>NS</sup> | -0,28<sup>NS</sup> | -0,83<sup>NS</sup> | -0,83<sup>NS</sup> | -0,78<sup>NS</sup> |
| *S. dampfi* | 0,90<sup>NS</sup> | -0,86<sup>NS</sup> | -0,58<sup>NS</sup> | -0,02<sup>NS</sup> | -0,86<sup>NS</sup> | -0,81<sup>NS</sup> | -0,85<sup>NS</sup> |
| *S. pedrosae* | -0,04<sup>NS</sup> | 0,36<sup>NS</sup> | 0,45<sup>NS</sup> | 0,69<sup>NS</sup> | 0,25<sup>NS</sup> | 0,32<sup>NS</sup> | 0,20<sup>NS</sup> |
| *X. affinis* | -0,81<sup>NS</sup> | 0,62<sup>NS</sup> | 0,26<sup>NS</sup> | -0,15<sup>NS</sup> | 0,69<sup>NS</sup> | 0,63<sup>NS</sup> | 0,71<sup>NS</sup> |

Tmax = maximum temperature, Tmin = minimum temperature, Tmed = average temperature, Umax = maximum humidity, Umin = minimum humidity, Umed = average humidity, P (mm) = precipitation. \* significant at 5% probability by Pearson's correlation; NS = not significant.

In Environment 1, there was no significant correlation for any of the climatic variables analysed in relation to species abundance in the four sampling periods. The species *Premnobius cavipennis* showed a high negative correlation for the minimum temperature variable in Environments 2 and 3, which means that with decreasing temperatures, the number of individuals increased.

Similarity analysis of the three environments and the diversity of the Scolytinae (Fig. 2) demonstrates the formation of a group of greater similarity between Environments 1 and 3 and their dissimilarity with Environment 2.

**Fig 2.**
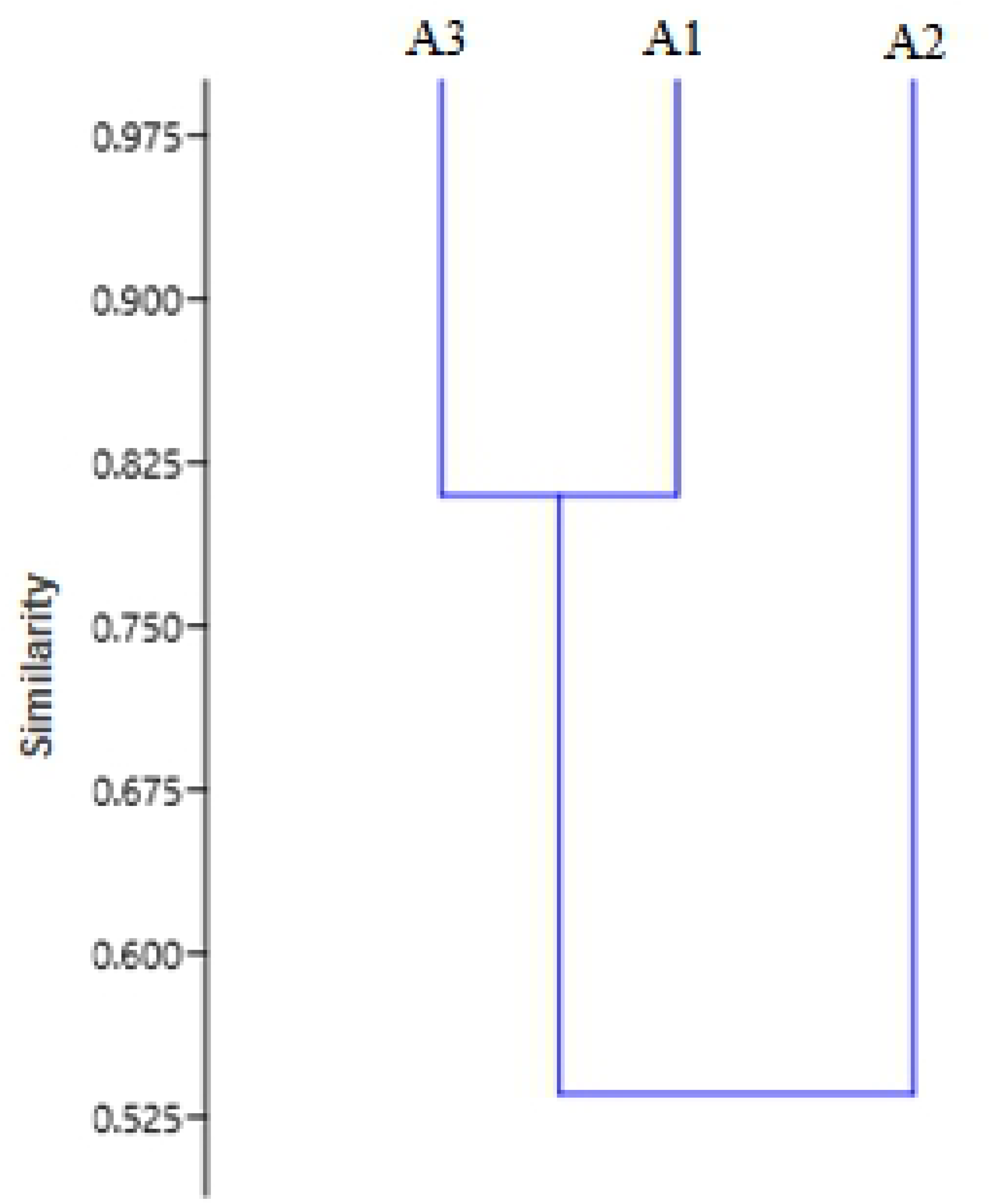
Similarity dendrogram (co-phenetic coefficient = 0.98) for species of Scolytinae collected in different environments in four sampling periods in the Southern Amazon.

## Discussion

The results of this study show a wide diversity of Scolytinae associated with native or homogeneous Brazil nut stands. Despite different vegetative compositions, the three environments had a wealth of similar species, with the largest number of species seen in Environment 3, which showed a greater diversity of plant species, greater conservation and concomitant and greater dynamics of ecological niches; it also demonstrated that the heterogeneity of the environments influenced the richness of the species of this group of wood-boring insects.

A significant number of species was also identified in Environment 2, which presented two species less than Environment 3 and 1 species more than Environment 1. The wealth of Scolytinae observed in Environment 2 suggests their effective association with the conditions of the homogeneous production of *Bertholletia excelsa* and, in addition, indicates the influence of high planting density, abundance of woody material and volatile substances, as well as decomposing plant material, such as leaves and branches of different sizes of Brazil nut trees found throughout the area during the sampling periods, on the occurrence of Scolytinae in the environment. The accumulation of litter in dense forest environments positively interferes with the development of beetles [24]. In addition, in homogeneous environments, phytophagous insects can easily adapt and colonise, becoming pests, due to the excessive supply of food [25].

Muller and Andreive [26] obtained similar results in the number of species collected among the different studied forest formations, where they found 28 species in the area of unaltered dense rainforest, 29 species in the area of altered dense rainforest and 21 species in the planting area homogeneous of *Eucalyptus grandis* (Hill ex Maiden). Rodriguez, Cognato and Righi [27], in a study of the diversity of Scolytinae in native forests and intercropped and homogeneous forest systems, also found results similar to those of the present study, with little variation between the richness of species in the environments, especially in the anthropized ones, observing 19 species for a native forest fragment, 24 species in the area of agroforestry system and 22 species in a homogeneous rubber plantation (*Hevea brasiliensis* Willd. Ex A. Juss. Müll. Arg.).

It is important to point out, as shown in the Venn Diagram (Fig 1), that in Environment 2, there were 16 species common to native environments, which corroborates that several species of Scolytinae found favourable conditions for their development in the homogeneous systems. The species that were restricted to some environments presented low frequency values in relation to the sampled totals and therefore were not classified, in the entomofaunistic analysis, as possible ecological indicators. Species with low population levels, categorised as non-dominant, infrequent, rare or dispersed, should be considered important in assessing the environment, since these species, under certain conditions of resource availability or interspecific relationships, can raise their fauna levels [28].

The majority of species collected in this study fall into the group of xylomycetophages, which are highly common in tropical environments, with the exception of the species *Cnesinus dryografus*, phloem borer, and *Coccotrypes palmarum*, seed pest.

The Shannon-Wiener Diversity Index was lower in Environment 1 (1.53), and for Environment 2 (1.75) and Environment 3 (1.71), the values were similar. In general, structurally more complex environments, which present greater dynamics of ecological niches, such as native systems, are characterised by greater diversity. However, it is noteworthy that it is not uncommon in studies of Scolytinae diversity to verify rates for homogeneous or intercropped areas, similar or even higher, to those calculated for native areas [24, 27, 29]. Scolytinae mainly act in the degradation of plant material that accumulates in environments, their diversity being related, among other environmental factors, to the amount of litter. Forested areas generally have a greater amount of material that can serve as a place of development, such as plant residues, broken trunks and branches, which provide conditions for the population growth of Scolytinae [30]. Thus, Environment 2, due to the high density of Brazil nut trees and the great availability and deposition of plant material, provided conditions for an expressive number of species and an abundance of Scolytinae individuals, which is also reflected in the value of the Shannon-Wiener index, which was similar to that of Environment 3. In addition, it is also possible to infer that the small difference found between the index values between these two environments may have been determined by equitability, which was slightly higher for Environment 2.

The equitability of Environment 2 (0.55) and Environment 3 (0.52) also showed similar values, indicating greater uniformity in the distribution of individuals between species compared to Environment 1, which obtained the lowest value (0.48). The values of the Shannon-Wiener diversity and the Equitability index in the three environments were lower than those observed in the evaluation of Scolytinae in a production system of *Eucalyptus urophylla* x *Eucalyptus grandis* in northern Brazil, which obtained the value of 2 for this index. And an Equitability index value of 0.80 [31].

The highest value of the Simpson diversity index was found for Environment 2 (0.72), most likely because this index is directly related to the dominance of certain species in the evaluated community, according to the principles established by Simpson [32]. This environment presented the largest number of species classified as dominant, which are considered as those that receive the impact of the environment and change it [33]; therefore, these species have significant importance for population monitoring, since they can cause the appearance or disappearance of other species.

The species *Xyleborus affinis* was classified as super-dominant in the three studied environments, with frequency values above 40% in relation to the total number of individuals in each environment. This species is xylomycetophagous and common in almost all types of forest environments; it can cause damage to the wood by boring or even staining woody fabrics, due to the presence of a symbiotic fungus [34]. *Xyleborus affinis* was also highly representative in studies evaluating wood-boring insects (Coleoptera) in savanna areas in southern Mato Grosso [29, 35, 36], although with lower frequency values than those found in this study.

The species *Xyleborus ferrugineus* was dominant in Environments 1 and 2. It is common in forest environments [34], and in addition to directly causing damage, it is also a vector of the pathogenic fungus *Ceratocystis fimbriata* [37].

The species classified as indicators allow characterising environments in which ecological changes occurred. In this sense, species of the genus *Cryptocarenus* are indicators of environments in a state of ecological disturbance [16]. The species *Premnobius cavipennis* has been reported in several studies in homogeneous forest systems and in altered forest remnants [14, 26, 29, 38, 39]. *Xyleborus spinulosus* was verified as a dominant species in homogeneous eucalyptus systems [29, 31, 39]. In general, the species of the genera *Sampsonius* have been recorded in altered native areas or forest stands, albeit at low frequencies. The species *X. affinis* and *X. ferrugineus*, which were also classified as indicators, can be classified as generalist species given their wide distribution in different phytographic regions in Brazil, being common species in almost all forest environments in the typologies of Mato Grosso [34, 40] and with records of economic damage in homogeneous production systems

The indicator species of the three environments, analysed in the four sampling periods, showed variations in their fauna categories throughout the year, with the exception of *Xyleborus affinis*, which maintained the same fauna pattern regardless of the environment of occurrence. However, correlation analysis showed that only the species *P. cavipenis* was significantly negatively correlated with the minimum temperature in Environments 2 and 3, that is, as there was a reduction in the minimum temperature, more individuals were sampled, suggesting that this species has a greater preference for environments with milder temperatures.

Different results were verified in the evaluation of Scolytinae in the production system of *Eucalyptus urophylla* x *Eucalyptus grandis* in Southern Amazonia [31]. The authors found that the species *Cryptocarenus diadematus* correlated negatively with the maximum and average temperatures and positively with the minimum relative humidity and with rainfall. The species *Cryptocarenus seriatus* showed a positive correlation with the maximum temperature and *Cryptocarenus hevea* with precipitation. Monteiro, Garlet and Carvalho [41] studied the occurrence of Scolytinae in a consortium of Brazil nut and rubber tree, also in the Southern Amazon, and found that the species *Cryptocarenus diadematus* was negatively correlated with the average and maximum temperatures and positively with the minimum temperature, maximum humidity, minimum humidity and precipitation. In the same study, these authors found a positive correlation between precipitation and abundance of *Cryptocarenus seriatus*.

Similar to the results of this study, no significant correlation was observed between the climatic variables and the population fluctuations of *Xyleborus affinis*, *Sampsonius dampfi* and *Premnobius cavipennis* in a study of the occurrence of wood-boring beetles in the savanna in southern Mato Grosso [35]. Machado and Costa [42] did not observe a significant correlation between climatic variables and the species of Scolytinae identified in their study.

The climatic variables are directly related to the flight, reproduction and dispersion of these insects [34]; in addition, they influence the physiological conditions of the trees, making them more or less susceptible to interaction with broachers. Sampling with shorter intervals may allow a better understanding of environmental factors and the population dynamics of Scolytinae. In addition, it must be considered that several interactions also occur, given the existence of microclimate relationships produced in forest environments, as these can promote microclimates in divergence from the regional climate, where variables such as wind, temperature, humidity and rainfall can change due to composition and vegetation levels, as well as allowing the existence of distinct niches for several species [43].

In the present study, the non-significance of most of the analysed climatic variables may be related to the small variation of these over the months analysed. The population fluctuation of Scolytidae varies among seasons and is correlated with the life cycle, biological opportunity and other environmental factors [44]. Furthermore, monitoring the population fluctuation of Scolytinae throughout the year is essential for the construction of strategic bases for integrated pest management in production systems of Brazil nut.

The grouping of similarities (Fig. 2) verified between the three environments can be understood by the structural complexity of the vegetation and management impact in each analysed environment. The environments that are most similarly grouped (1 and 3) are environments with a similar vegetation composition, although in part of Environment 1, pastures occur. In contrast, Environment 2 has a homogeneous structure of individuals of *Bertholletia excelsa* and greater structural simplicity, favouring the occurrence of an expressive number of species of Scolytinae. It can therefore be said that the structural complexity of vegetation and environmental disturbances are the main factors that contribute to Scolytinae diversity patterns [27,45].

Similar to the results found in this work, Rodriguez, Cognato and Righi [27], studying the diversity of Scolytinae in native forest areas as well as intercropped and homogeneous forest systems, found that the native forest environments were grouped in greater similarity, and these in dissimilarity in relation to each other to another grouping between homogeneous planting of rubber (*Hevea brasiliensis* Willd. ex A. Juss., Müll. Arg) and coffee cultivation (*Coffea arabica* L).

## Conclusions

The three Brazil nut areas analysed, with their differences in vegetation structure, presented similar species richness. In addition, of the 31 identified species, 16 were common to the three environments. Environment 2 presented an expressive number of Scolytinae species, which demonstrates their effective association with the silvicultural conditions of the homogeneous production system.

The species *Xyleborus affinis* was the most representative in the three study environments and remained as a super-dominant species in all four sampled periods. *Xyleborus affinis*, *Cryptocarenus seriatus*, *Cryptocarenus diadematus* and *Premnobius cavipennis* represented 70% of the total of sampled individuals and should be considered as potential pest insects in production systems of Brazil nuts.

This shows that population monitoring of Scolytinae in Brazil nut production systems, especially homogeneous systems, is of fundamental importance for the construction of strategies for integrated pest management and the sustainability of these agroecosystems in the Amazon.

## Acknowledgments

We thank Prof. Dr. Eli Nunes Marques for his collaboration in species identification.

## Author Contributions

Conceptualisation: Marcus Henrique Martins e Silva, Juliana Garlet.

Formal Analysis: Marcus Henrique Martins e Silva, Juliana Garlet.

Investigation: Marcus Henrique Martins e Silva, Fernando Luiz Silva, Carla da Silva Paula. Methodology: Marcus Henrique Martins e Silva, Juliana Garlet.

Project Administration: Marcus Henrique Martins e Silva, Juliana Garlet.

Supervision: Juliana Garlet.

Visualisation: Marcus Henrique Martins e Silva.

Writing – Original Draft Preparation: Marcus Henrique Martins e Silva, Juliana Garlet.

Writing – Review & Editing: Marcus Henrique Martins e Silva, Juliana Garlet.

